# An evolutionary medicine perspective on Neandertal extinction

**DOI:** 10.1101/047209

**Authors:** Alexis P. Sullivan, Marc de Manuel, Tomas Marques-Bonet, George H. Perry

**Author notes:** Corresponding Author: George H. Perry.

## Abstract

The Eurasian sympatry of Neandertals and anatomically modern humans – beginning at least 45,000 years ago and lasting for more than 5,000 years – has long sparked anthropological interest into the factors that potentially contributed to Neandertal extinction. Among many different hypotheses, the “differential pathogen resistance” extinction model posits that Neandertals were disproportionately affected by exposure to novel infectious diseases that were transmitted during the period of spatiotemporal sympatry with modern humans. Comparisons of new archaic hominin paleogenome sequences with modern human genomes have confirmed a history of genetic admixture – and thus direct contact – between humans and Neandertals. Analyses of these data have also shown that Neandertal nuclear genome genetic diversity was likely considerably lower than that of the Eurasian anatomically modern humans with whom they came into contact, perhaps leaving Neandertal innate immune systems relatively more susceptible to novel pathogens. In this study, we compared levels of genetic diversity in genes for which genetic variation is hypothesized to benefit pathogen defense among Neandertals and African, European, and Asian modern humans, using available exome sequencing data (six chromosomes per population). We observed that Neandertals had only 31-39% as many nonsynonymous (amino acid changing) polymorphisms across 73 innate immune system genes compared to modern human populations. We also found that Neandertal genetic diversity was relatively low in an unbiased set of balancing selection candidate genes for primates – genes with the highest 1% genetic diversity genome-wide in non-human apes. In contrast, Neandertals had similar to higher levels of genetic diversity than humans in 13 major histocompatibility complex (*MHC*) genes. Thus, while Neandertals may have been relatively more susceptible to some novel pathogens and differential pathogen resistance could be considered as one potential contributing factor in their extinction, this model does have limitations.

## Introduction

Following their divergence from the modern human lineage ~550 kya^1–3^, Neandertals (*Homo neanderthalensis*) inhabited Eurasia from at least 300 kya^4–6^ to ~41-39 kya^7^. In contrast, anatomically modern humans very likely evolved in sub-Saharan Africa, where the earliest recognized skeletal remains date to 155 kya^8,9^. Modern humans then expanded out of Africa and spread from the Middle East into Europe by at least 45 kya^10–12^ and Asia by 80 kya^13,14^. Within 6 kya after modern human appearance in Europe, Neandertals no longer appear in the fossil record^15^.

There are numerous hypotheses regarding Neandertal extinction. For example, some scholars have suggested that climate fluctuations between 40-30 kya played a key role in this process^16–18^. However, this hypothesis has been discounted on the basis of the simultaneous Eurasian presence of anatomically modern humans for most of the 40-30 kya period combined with the *in situ* established environmental resilience of Neandertals^19^. That is, if anything, modern humans would more likely have been adversely affected by this environment than Neandertals, given the survival and likely adaptation of the latter to the harsh Eurasian climate for hundreds of thousands of years^20^.

Most other extinction hypotheses focus on Neandertal-modern human competition^21–25^. For example, potentially shorter inter-birth intervals for modern humans could have allowed more rapid population growth compared to Neandertals, facilitating eventual replacement^26,27^. Alternatively, anthropologists have speculated that the intelligence and language capabilities of anatomically modern humans were greater than those of Neandertals^28–31^, perhaps facilitating competitive hunting and other advantages such as through the development of more efficient tool technologies^32^. A recent proposal is that modern humans benefitted from the early domestication of dogs, who may have aided large animal hunts to increase caloric yields for the modern humans and fuel their rapid population growth and ultimately larger sizes^33^.

Thinking of the important role of disease in population dynamics, Wolff and Greenwood^34^ suggested that viral disease transmission from modern humans to Neandertals could have contributed to the ultimate disappearance of the latter. Houldcroft and Underdown^35^ recently echoed and expanded on these notions. This “differential pathogen resistance” model would require an anatomically modern human pathogen (or pathogens) of limited virulence for out-of-Africa migrating human populations, but that would have strongly affected immunologically naïve Neandertal populations upon contact and transmission. Moreover, the viability of this scenario also requires i) relatively fewer Neandertal pathogens/ strains at the time of contact, ii) some mechanism by which modern humans might not have been as negatively affected as Neandertals upon novel pathogen exposure, or iii) both of these factors to be present. In this paper we assess the plausibility of the differential pathogen resistance model by comparing levels of genetic diversity in genes for which genetic variation is hypothesized to benefit pathogen defense between Neandertal and modern human populations.

Recent advances in genomic sequencing technologies and ancient DNA methods have facilitated the generation of a high-quality Altai Neandertal nuclear genome sequence from Siberia (dated to ~50,000 ya)^36^. When analyzed in combination with modern human genomic data, this genome has provided convincing evidence that anatomically modern humans and Neandertals interbred^3,36–39^. The requisite intercourse demonstrates at least some level of direct contact between these populations, and thus opportunities for the transfer of infectious diseases.

Moreover, analyses of both the high-coverage diploid nuclear genome sequence from the Altai Neandertal and mitochondrial DNA sequence data that are available for multiple Neandertal individuals suggest that Neandertal genetic diversity was substantially lower than that observed within modern human populations^3,36,40,41^. Recently, Castellano et al.^42^ used a DNA capture method to sequence the nuclear genome’s protein-coding regions – the ‘exome’ – from each of two additional Neandertals: individuals who lived ~49,000 ya^43^ and ~44,000 ya in Spain and Croatia, respectively. Observed levels of heterozygosity for these two Neandertals are also relatively low, suggesting that low nuclear genome genetic diversity was a general Neandertal characteristic rather than restricted to an Altai Neandertal population isolate^42^. Specifically, considering only sites with sequence coverage sufficient for single nucleotide polymorphism (SNP) identification for each of the three Neandertals, Castellano et al.^42^ observed only 30.3%, 44.9%, and 45.3% as many synonymous (i.e., not amino acid-changing) SNPs in the Neandertals compared to equal-sized population samples of modern human Africans, Europeans, and Asians, respectively.

Within genes directly related to immune function, greater functional genetic diversity increases the potential responsiveness of the immune system to foreign pathogens^34,44^. Balancing selection is thought to maintain advantageous functional diversity (i.e., nonsynonymous SNPs) within these genes^45,46^, and individuals with more genetic diversity across the genome tend to have higher fitness^44^. Based on population genetic theory, genetic drift is a relatively stronger force, while natural selection is relatively less effective, in smaller populations^47,48^. Thus, compared to a larger population, a population with a historically small effective population size may have lower genetic diversity in general across the genome *and* different patterns of diversity at loci affecting individual health and fitness. Indeed, along with relatively reduced overall genetic diversity, Castellano et al.^42^ observed a higher proportion of predicted damaging than benign nonsynonymous (amino acid-changing) SNPs in Neandertals compared to modern humans, consistent with the reduced effectiveness of purifying selection to remove or reduce the frequencies of strongly deleterious variants in Neandertals^49–53^.

In addition to purifying selection, other types of natural selection, including balancing selection, are likewise expected to be less effective in smaller populations. Thus, the generally low genetic diversity of Neandertals relative to humans may even be exacerbated at functional sites in genes related to immune function that would otherwise be preserved via balancing selection. Theoretically, such a difference could have facilitated the differential morbidity following contact and infectious disease transfer between Neandertals and modern humans potentially required under the Pleistocene epidemiological scenarios (the differential pathogen resistance model) of Wolff and Greenwood^34^ and Houldcroft and Underdown^35^.

In this study we specifically compared the levels and patterns of genetic variation between Neandertal and modern human populations at i) 73 genes associated with innate immune functions, ii) 164 virus-interacting protein genes, iii) 13 major histocompatibility complex (*MHC*) genes, and iv) the 1% of genes across the genome with the consistently highest levels of genetic diversity among four non-human ape species. Relatively lower genetic variation at these loci in Neandertal populations would be consistent with the differential pathogen resistance hypothesis that a greater susceptibility to novel pathogens relative to the modern humans with which they interacted could have been one contributing factor in the extinction of Neandertals. In contrast, similar levels of Neandertal and modern human genetic diversity would raise major questions about the plausibility of this epidemiological extinction hypothesis.

## Results

Castellano et al.^42^ produced a database of Neandertal and African, European, and Asian human exonic SNP genotypes based on the paleogenomic data for the three Neandertal individuals described above and high-coverage genome sequencing data for three human individuals from each of three continental regions. Ancestral and derived alleles were determined via comparisons to the gorilla and orangutan genome sequences^42^. Genotypes were estimated for only those positions covered by a minimum of six independent sequencing reads; the authors also tested higher minimum coverage cutoffs (up to 20x) that did not significantly affect the observed patterns of Neandertal versus modern human genetic diversity, so the lower coverage cutoff was used to include a larger number of sites in the analysis^42^. For our analysis, we considered only the autosomal sites with sufficient coverage for SNP genotyping among all individuals, so that the numbers of identified SNPs per population and their allele frequency distributions could be compared directly among the Neandertal and modern human African, European, and Asian samples (n = 6 chromosomes per population) as measures of genetic diversity.

### Patterns of genetic diversity and the effectiveness of selection

Following Castellano et al.^42^, we used the program PolyPhen-2^54^ to infer whether each nucleotide variant was nonsynonymous (amino acid changing; potentially functional) or synonymous (resulting in no change to the amino acid sequence; typically neutral with respect to fitness), and for the nonsynonymous mutations a “damaging” or “benign” prediction of their effects on protein structure and function. On the overall dataset, we observed patterns of genetic diversity that were similar to the primary findings of Castellano et al.^42^. Specifically, Neandertals had relatively fewer total SNPs than any of the modern human populations; this was true for both nonsynonymous and synonymous SNPs (**Figure 1; Supplemental Table 1**). In addition, Neandertals also had a higher proportion of nonsynonymous to total (nonsynonymous + synonymous) SNPs compared to the human populations (Neandertals = 0.500; human populations = 0.440-0.452; Fisher’s Exact Test for the Neandertal vs. European comparison; *P* = 2.29x10^−10^; see **Supplemental Table 1** for all comparisons) and a considerably higher proportion of predicted damaging to total nonsynonymous SNPs (Neandertal = 0.435; human = 0.291-0.297; *P* < 2.2x10^-16^ for the Neandertal-European comparison; **Supplemental Table 2**). Together, these results are consistent with the notion that purifying natural selection against potentially damaging nonsynonymous SNPs was relatively less effective in smaller Neandertal populations.

**Figure 1.**
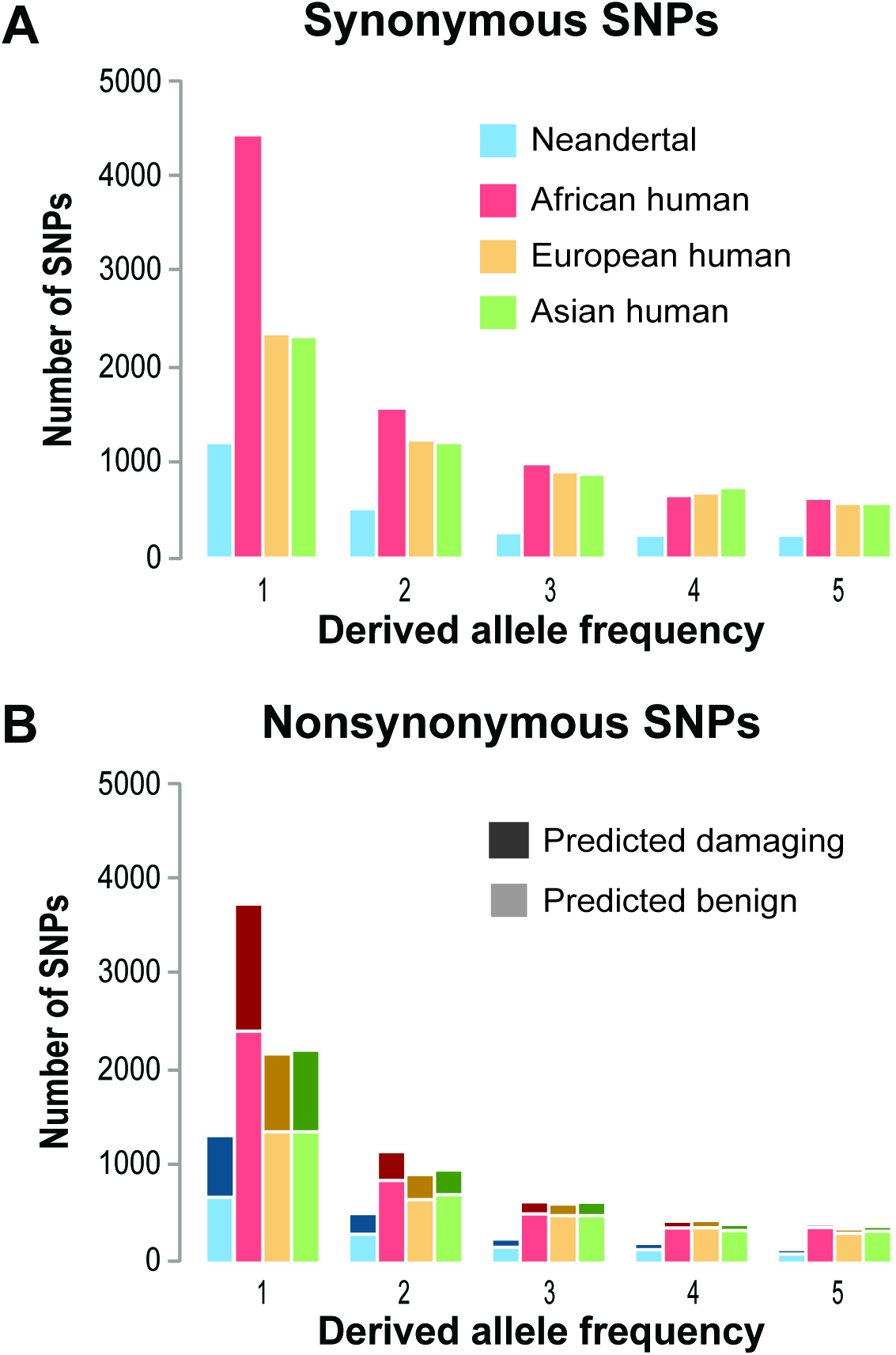
Genome-wide SNP derived allele frequency distributions for Neandertals and three modern human populations. The number of SNPs observed for each population at exome sites with sufficient sequence coverage for SNP identification in all individuals in the study, binned by derived allele frequency (of n = 6 chromosomes per population). (A) Synonymous SNPs. (B) Nonsynonymous SNPs, with “predicted benign” and “predicted damaging” SNPs separately indicated, based on PolyPhen-2^54^ estimates.

### Innate immune system, virus-interacting protein, and MHC gene diversity comparisons

To assess the plausibility of the differential pathogen resistance model, we next performed analyses focused on patterns of genetic diversity within two subsets of genes for which genetic diversity itself is thought to play an important role in pathogen defense-related immune functions. The first set was comprised of 73 innate immune receptor, signaling adaptor molecule, and complement pathway genes (“Innate Immune System Genes”; **Supplemental Table 3**), for example the toll-like receptor^55–57^ and mannose-associated serine protease genes^58^. Many of the proteins encoded by these genes are involved in the innate immune system’s first line of defense against diverse external microorganisms. The second set was comprised of 164 genes encoding for virus-interacting proteins that also have known antiviral activity or broader roles in the immune system^59^ (**Supplemental Table 4**). The second set was comprised of 13 *MHC* genes (**Supplemental Table 5**), critical immune system loci with among the strongest evidence for long-term balancing selection and long-term maintenance of allelic diversity in vertebrate genomes^60–63^. The patterns of Neandertal and modern human genetic diversity for these two gene sets were compared to those for the remaining 11,086 genes in our genome-wide dataset.

Neandertal and human nonsynonymous genetic diversity was similar between the genome-wide and innate immune system gene sets, with fewer nonsynonymous SNPs in Neandertals compared to any human population in both cases (**Figure 2A-D**). In fact, the relative number of Neandertal vs. human nonsynonymous SNPs is even slightly higher for the genome-wide set than for the innate immune system genes; e.g., with 54.1% as many Neandertal as European human nonsyonymous SNPs in the genome-wide set compared to only 39.0% for the innate immune genes. The direction of this result is consistent with expectations under a model of relatively reduced balancing selection effectiveness in Neandertals (i.e., if innate immune gene functional genetic diversity confers a fitness advantage). However, the observed Neandertal-human difference between the two gene sets is not significant based on Fisher’s Exact Tests (*P* = 0.33 for Neandertals-Europeans; see **Supplemental Table 6** for all comparisons). Based on the results of a permutation analysis with 10,000 sets of 73 random genes from the genome-wide set, the proportion of the number of Neandertal-human nonsynonymous SNPs is also not significantly lower than expected by chance (*P* = 0.1944 for Neandertals-Europeans; see **Supplemental Figure 1** for all comparisons). Regardless, our results do not provide any support for the possibility that strong balancing selection has maintained similar levels of nonsynonymous diversity in Neandertal and human innate immune system genes despite the lower effective population size and genome-wide diversity of Neandertals.

**Figure 2.**
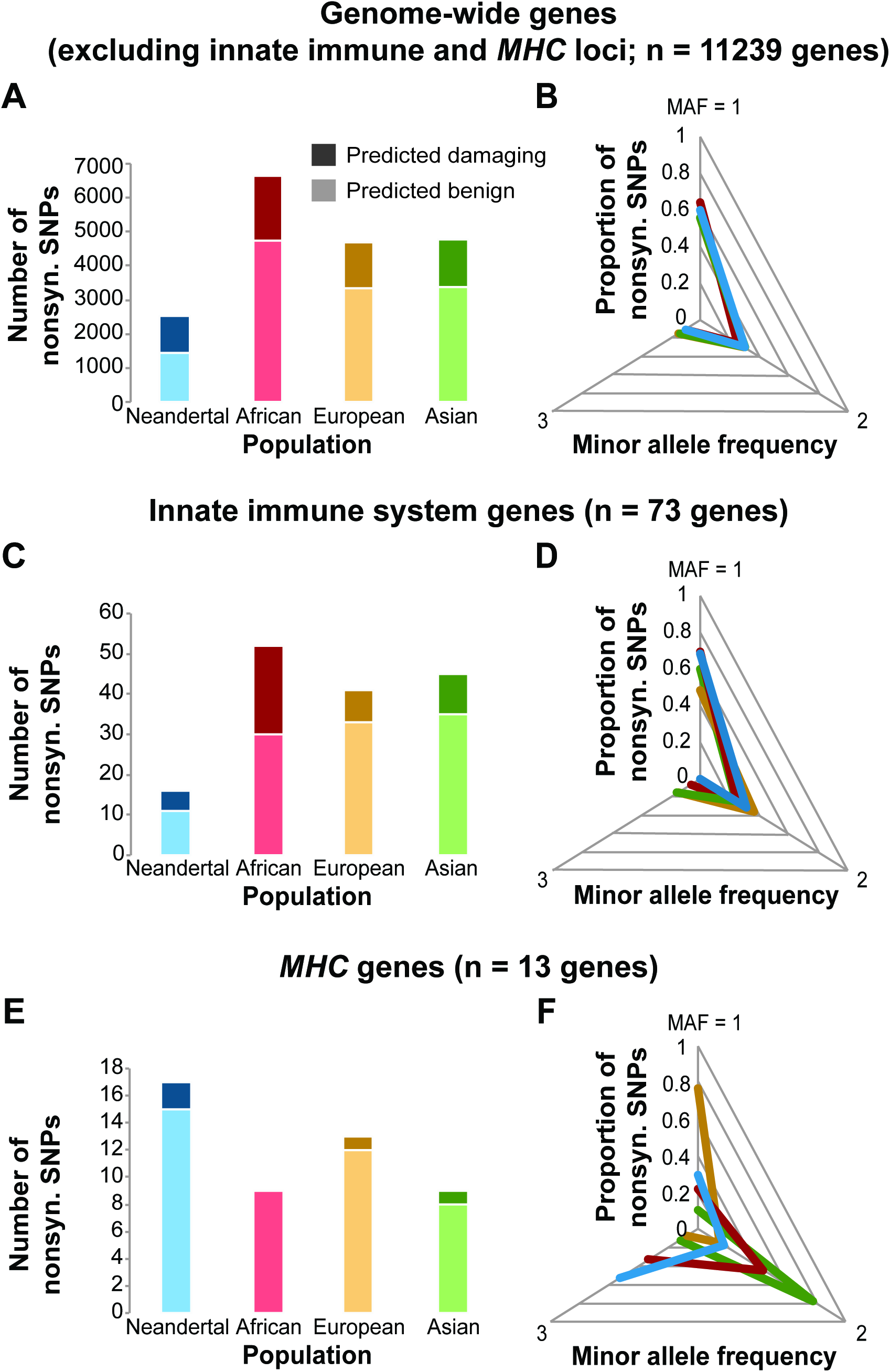
Patterns of Neandertal and modern human nonsynonymous SNP diversity genome wide, in innate immune genes, and in *MHC* genes. The number of nonsynonymous SNPs per population for sites with sufficient sequence coverage for SNP identification in all individuals in the study, with PolyPhen-2 “predicted benign” and “predicted damaging” SNPs indicated separately, and the proportion of the nonsynonymous SNPs with minor allele frequencies = 1, 2, and 3 (of n = 6 total chromosomes per population). (A-B) Nonsynonymous SNPs in the genome-wide set of n = 11,086 genes that excludes innate immune receptor, signaling adaptor molecule, complement pathway, virus-interacting protein, and *MHC* genes. (C-D) Nonsynonymous SNPs in the set of n = 73 innate immune receptor, signaling adaptor molecule, and complement pathway genes. (E-F) Nonsynonymous SNPs in the set of n = 164 virus-interacting protein genes with known antiviral or immune activities. (G-H) Nonsynonymous SNPs in the set of n = 13 *MHC* genes.

The relative number of Neandertal vs. human nonsynonymous SNPs observed among the virus-interacting protein genes was also low (**Figure 2E-F**), in this case at similar proportions to the genome-wide gene set (**Supplemental Table 7; Supplemental Figure 2**).

In contrast, there was a greater number of total nonsynonymous SNPs observed across the 13 *MHC* genes for Neandertals (17) than for any of the three human populations (9, 13, and 9 for the African, European, and Asian population samples, respectively; **Figure 2G**). Compared to both the genome-wide set of genes and the innate immune system genes, the number of Neandertal relative to human *MHC* gene nonsynonymous SNPs is significantly greater than expected by chance when assessed with Fisher’s Exact Tests (e.g., Neandertals-Europeans for *MHC* vs. genome-wide set; *P* = 0.02; see **Supplemental Table 8** for all comparisons) and, for some but not all population comparisons, with permutation analyses (*P* = 0.1556 for Neandertals-Europeans; *P* = 0.0759 for Neandertals-Asians; *P* = 0.0222 for Neandertals-Africans; **Supplemental Figure 3**).

Additionally, we observed 9/17 (52.9%) of the Neandertal *MHC* nonsynonymous SNPs at intermediate frequency (i.e., with the minor allele observed on 3 out of the 6 chromosomes in the population; **Figure 2H**), a significantly higher proportion than observed for Neandertal nonsynonymous SNPs in the genome-wide, non-immune gene set (260/2527; 10.3%; Fisher’s Exact Test; *P* = 1.56x10^−5^; **Supplemental Table 9**). The proportion of Neandertal *MHC* intermediate frequency nonsynonymous SNPs is also relatively higher for Neandertals than for any of the human populations, although with the small sample sizes not all comparisons were statistically significant (Fisher’s Exact Tests; *P* = 0.43 for Neandertals-Africans; *P* = 0.02 for Neandertals-Europeans; *P* = 0.09 for Neandertals-Asians; **Supplemental Table 10**). The Neandertal-human *MHC* genetic diversity patterns remained similar after we excluded one random individual from each human population and the Altai Neandertal individual, whose genomic *MHC* region may harbor some DNA segments introgressed from humans^64^ (**Supplemental Figure 4**).

In combination, the relatively large number of Neandertal *MHC* nonsynonymous SNPs and the high proportion of those variants observed at intermediate frequencies suggest that at least for these critical immune loci, the heterozygous fitness advantage for Neandertals was sufficiently strong to offset the lower effective size of this population, leading to similar or even higher putatively functional diversity compared to modern human populations.

### Balancing selection candidate gene comparisons

Finally, we sought to compare patterns of Neandertal-human genetic diversity across genes for which evidence of balancing selection has been identified without respect to gene function. However, were we to analyze candidate genes detected using human population genomic data, we would introduce bias into our Neandertal vs. human comparison; thus we cannot use gene lists from the majority of genome-wide analyses of balancing selection in mammals that have been published to date^46,60,63,65–68^.

Instead, we analyzed genome-wide sequence data from 55 total individuals from population samples of four non-human ape species, chimpanzee (*Pan troglodytes ellioti*), bonobo (*Pan paniscus*), gorilla (*Gorilla gorilla gorilla*), and orangutan (*Pongo abelii*)^69^, to identify an unbiased (with respect to our analysis) set of candidate balancing selection genes. Specifically, we estimated per-species nucleotide sequence diversity as the average proportion of pairwise differences (π) for the coding regions of each gene. There were 7,259 genome-wide genes with ≥ 500 “callable” sites across all four species (see *Methods*) for which there was also at least one variable site in our Neandertal-modern human genetic diversity database. Within each species, we computed π value percentiles for each gene, and then summed the per-gene percentile values across species. Using this approach, we identified the 73 (top 1%) genes with the consistently highest genetic diversity among these non-human ape species (“top 1% ape diversity genes”; **Supplemental Table 11**).

Relative to all 7,259 genes and following correction for multiple tests, the top 1% ape diversity set was significantly enriched for genes with immune system-related Gene Ontology functional categories including “MHC protein complex” (observed = 3 genes; expected = 0.09 genes; adjusted *P* = 0.0015), “positive regulation of leukocyte activation” (observed = 6 genes; expected = 0.62 genes; adjusted *P* = 0.0061), and “immune system process” (observed = 14 genes; expected = 4.92 genes; adjusted *P* = 0.0098). Full results from the Gene Ontology enrichment analysis are provided in **Supplemental Table 12** (see also **Supplemental Figure 5**). In addition to the *MHC* loci, the top 1% ape diversity set contains *OAS1*, another gene for which high allelic diversity appears likely to have been maintained across multiple species by long-term balancing selection^70^. Together, these features suggest that our top 1% ape diversity genes are likely at least enriched for those affected by balancing selection in hominoid primates. Thus, it is appropriate to compare Neandertal and modern human genetic diversity at these loci as an additional component of our assessment of the underlying mechanics of the differential pathogen resistance model for Neandertal extinction.

Across the top 1% ape diversity genes, we observed only 27%-58% as many nonsynonymous SNPs for Neandertals compared to the modern human populations (**Figure 3C**). The magnitude of this difference was similar to that for the remaining genome-wide genes (**Figure 3A**; Fisher’s Exact Test; *P* = 0.69 for Neandertals-Europeans; see **Supplemental Table 13** for all comparisons). Thus, the pattern of Neandertal vs. modern human nonsynonymous genetic diversity for the top 1% ape diversity gene set was more similar to that observed for the innate immune system genes than the *MHC* genes.

**Figure 3.**
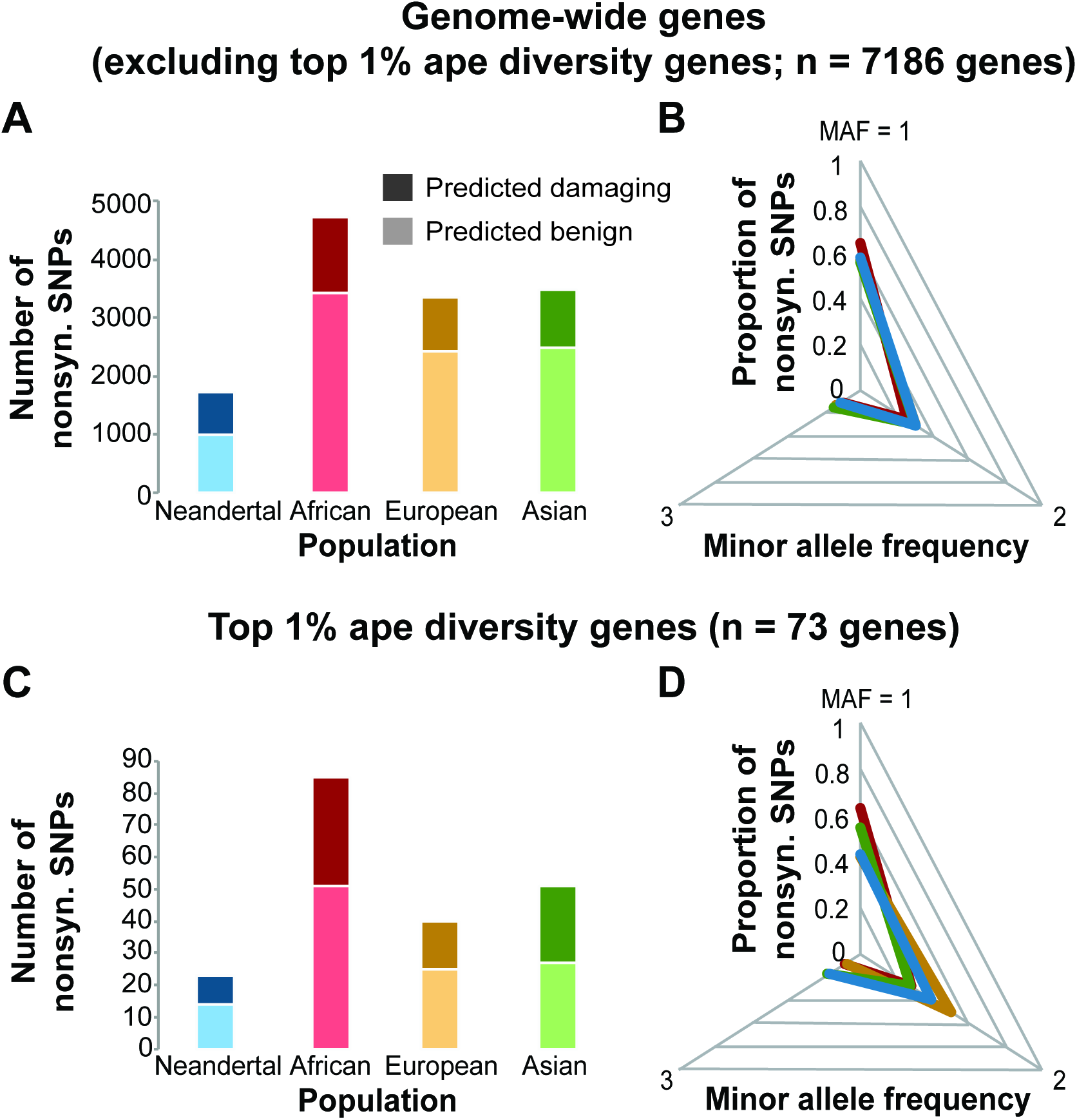
Patterns of Neandertal and modern human nonsynonymous SNP diversity genome wide and for genes with high genetic diversity in non-human apes. The number of nonsynonymous SNPs per population for sites with sufficient sequence coverage for SNP identification in all individuals in the study, with PolyPhen-2 “predicted benign” and “predicted damaging” SNPs indicated separately, and the proportion of the nonsynonymous SNPs with minor allele frequencies = 1, 2, and 3 (of n = 6 total chromosomes per population). (A-B) Nonsynonymous SNPs in the genome-wide set of n = 7,186 genes with at least 500 “callable” sites with sufficient coverage and mapping quality for SNP identification in a great ape population genomics panel (see *Methods*), excluding the top 1% highest diversity genes among four non-human ape species. (C-D) Nonsynonymous SNPs in the set of n = 73 genes with the consistently highest levels of genetic diversity among the four non-human ape species.

## Discussion

The differential pathogen resistance model of Neandertal extinction posits that after contact with anatomically modern humans who migrated Out of Africa, Neandertals may have been disproportionately affected by transferred infectious diseases. Our goal in this study was to assess the underlying plausibility of this model by testing whether Neandertal genetic diversity at critical immune system loci was significantly lower than that for the modern human populations with whom they came into contact – a possibility, given lower levels of general, genome-wide Neandertal genetic diversity^42,71,72^. The model would *not* require Neandertals to be depauperate of distinct and evolutionarily advantageous genetic variants at immune system loci. Indeed, evidence of the adaptive introgression of Neandertal toll-like receptor and *MHC* gene variants into modern human populations strongly suggests otherwise^73–75^. Rather, the mechanics of the differential pathogen resistance model could still be relevant if there were simply *fewer* such variants and substantially *lower* functional genetic diversity overall at immune system loci in Neandertals compared to sympatric modern human populations.

Are our results consistent with the differential pathogen resistance model for Neandertal extinction? Not in full. While Neandertal nonsyonymous genetic diversity at innate immune system, virus-interacting protein, and non-human ape high-diversity gene loci was indeed low compared to that observed for humans, Neandertal *MHC* diversity was similar or even higher than that for humans, in multiple respects. Thus, future models that incorporate epidemiological mechanisms as contributing factors to Neandertal extinction should proceed with caution, as there are specific genes for which balancing selection in Neandertals appears to have overcome the lower effective size and lower levels of genome-wide genetic diversity in this population. That said, there still may have been an aggregate differential pathogen resistance effect from lower functional diversity across innate immune system and other critical pathogen defense genes. Looking forward, this hypothesis could be explored further with the combination of expanded Neandertal paleogenomic population data and broad experimental/ functional comparisons of Neandertal vs. modern human innate immune gene diversity.

## Materials and Methods

We downloaded the Neandertal exome DNA capture data published by Castellano et al.^42^ (http://cdna.eva.mpg.de/neandertal/exomes/VCF). Specifically, we considered the SNP genotype data for autosomal chromosomes from the “combined” VCF files from this dataset, in which SNP genotypes for each of the 13 individuals in the dataset (three modern humans of African descent, three modern humans of European descent, three modern humans of Asian descent, three Neandertal individuals, and one individual from the archaic hominin Denisovan population) were provided for only the individuals with a minimum of six independent sequencing reads at that position. There were a total of 69,230 autosomal SNPs in this dataset. We removed n = 32,416 SNPs for which genotypes were not estimated for all of the Neandertal and modern human individuals and n = 17 SNPs with more than 2 identified alleles (e.g., A/T/C variably present at one position).

The remaining 36,797 autosomal SNPs were submitted to PolyPhen-2’s online HumDiv server via batch query for identification of nonsynonymous and synonymous SNPs and estimates of predicted functional consequences for the nonsynonymous SNPs^42,54^. PolyPhen-2 classified each SNP as “missense” (nonsynonymous), “coding-synon” (synonymous), “nonsense” (stop codon), “utr-5” (5’ untranslated region), “utr-3” (3’ untranslated region), “intron” (non-coding variants). Our subsequent analysis focused on the nonsynonymous (n = 16,139) and synonymous (n = 18,095) SNPs. For each nonsynonymous SNP, PolyPhen-2 also provided “benign” (n = 10,558), “possibly damaging” (n = 2,222), and “probably damaging” (n = 3,317) predictions^54^. PolyPhen-2 failed to make functional predictions for a small number of nonsynonymous SNPs (n = 42), which were removed from our subsequent analyses. SNPs in the predicted possibly and probably damaging categories were combined into one “predicted damaging” category for our analyses. PolyPhen-2 assembled gene names from the UCSC knownGene transcripts/database (hg19/GRCh37), and these gene names were used for later identification. The annotated database of nonsynonymous and synonymous SNP genotypes and population frequencies that we analyzed for this study are available in **Supplemental Database 1** (https://scholarsphere.psu.edu/files/v405s943f). All analyses of the SNP genotype data were performed using the R statistical environment^76^.

In addition, to compare patterns of synonymous and nonsynonymous SNP derived allele frequencies among populations (e.g., see **Figure 1**) we determined the derived and ancestral states for the alleles of each SNP by comparison to the orthologous chimpanzee, gorilla, and orangutan nucleotides based on alignments to reference genomes for those species as provided by Castellano et al.^42^. Ancestral alleles were classified as those that matched the orthologous nucleotides for all three non-human species. SNPs with variability, missing data, or nucleotides different than either of the two Neandertal/human alleles at the orthologous positions among the three non-human species (n = 538 nonsynonymous and n = 738 synonymous SNPs) were not considered in our analyses of derived allele frequencies. Note that our comparisons of the patterns of genetic diversity among gene categories (e.g., see **Figures 2 and 3**) did not require derived allele frequency information; thus, the SNPs with variability, missing data, or nucleotides different than the two Neandertal/human alleles were still included in that analysis to avoid bias against highly variable loci (e.g., those with cross-species polymorphisms maintained by long-term balancing selection).

To identify the set of genes with consistently high levels of genetic diversity among non-human apes, we analyzed whole-genome genetic diversity data that were previously published by Prado-Martinez et al.^69^ for 87 great ape individuals. Briefly, in that study DNA from each individual was sequenced on an Illumina platform to ~25x sequence coverage. Reads were mapped to the human reference genome (hg18). Genotypes were estimated only for sites that met criteria for sequence coverage, base quality, and mapping quality (“callable sites”), as described^69^. For our genetic diversity analysis, we focused on a subset of these data (n = 55 individuals), considering the one population from each ape species or species group with the largest sample size: chimpanzee (*Pan troglodytes ellioti*; n = 10), bonobo (*Pan paniscus*; n = 13), gorilla (*Gorilla gorilla gorilla*; n = 27), and orangutan (*Pongo abelii*; n = 5). We obtained hg18 gene coding region coordinates from the knownCanonical gene database using the table browser at the UCSC Genome Bioinformatics Site (genome.ucsc.edu/index.html). For each gene also in the Neandertal-human genetic diversity database we estimated nucleotide diversity as the average proportion of pairwise differences (π) for the callable sites of the coding regions^77^. The great ape gene diversity data analyzed in this study are available in **Supplemental Database 2** (https://scholarsphere.psu.edu/files/5d86p0267). Gene Ontology enrichment analyses were performed using the WEB-based Gene SeT AnaLysis Toolkit (WebGestalt). Functional category enrichments were statistically evaluated with hypergeometric tests and p-values were adjusted for multiple tests using the method of Benjamini & Hochberg^78^.

## Acknowledgments

We thank Sergi Castellano and Luis Barreiro for their discussions on this project, and David Enard and Martin Kuhlwilm for helpful analytical suggestions. The computational resource instrumentation used in this study was funded by the National Science Foundation (OCI–0821527).

## Supplementary Materials

**Supplemental Database 1**: Database of nonsynonymous and synonymous SNP genotypes and population frequencies (https://scholarsphere.psu.edu/files/v405s943f)

**Supplemental Database 2**: Database of great ape gene diversity data (https://scholarsphere.psu.edu/files/5d86p0267)

**Supplemental Figure 1**: Innate immune system gene permutation analyses – 10,000 sets of 73 randomly selected genes containing nonsynonymous SNPs

**Supplemental Figure 2**: Virus-interacting protein gene permutation analyses – 10,000 sets of 164 randomly selected genes containing nonsynonymous SNPs

**Supplemental Figure 3**: *MHC* gene permutation analyses – 10,000 sets of 13 randomly selected genes containing nonsynonymous SNPs

**Supplemental Figure 4**: Patterns of Neandertal and modern human nonsynonymous SNP diversity in *MHC* genes (n = 13) excluding the Altai Neandertal and one random modern human per population

**Supplemental Figure 5**: Significantly enriched gene ontology categories (red) among top 1% ape diversity genes

**Supplemental Table 1**: A comparison of genome-wide nonsynonymous SNPs versus total (nonsynonymous + synonymous) SNPs between Neandertal and modern human populations

**Supplemental Table 2**: PolyPhen-2 predictions for genome-wide nonsynonymous SNPs – damaging versus not damaging – for Neandertal and modern human populations

**Supplemental Table 3**: List of Innate immune system genes

**Supplemental Table 4**: List of virus-interacting protein genes with known antiviral or broader immune activities

**Supplemental Table 5**: List of *MHC* genes

**Supplemental Table 6**: A comparison of nonsynonymous SNPs (benign + damaging) in innate immune system genes versus genome-wide genes (not including innate immune genes) between Neandertal and modern human populations

**Supplemental Table 7**: A comparison of nonsynonymous SNPs (benign + damaging) in virus-interacting protein genes versus genome-wide genes (not including virus-interacting protein genes) between Neandertal and modern human populations

**Supplemental Table 8**: A comparison of nonsynonymous SNPs (benign + damaging) in *MHC* genes versus genome-wide genes (not including *MHC* genes) between Neandertal and modern human populations

**Supplemental Table 9**: Neandertal Minor Allele Frequency (MAF) comparison of nonsynonymous SNPs in *MHC* genes

**Supplemental Table 10**: A comparison of nonsynonymous SNP (benign + damaging) Minor Allele Frequencies (MAFs) in *MHC* genes between Neandertal and modern human populations

**Supplemental Table 11**: List of top 1% ape diversity genes

**Supplemental Table 12**: Results of the top 1% ape diversity genes Gene Ontology enrichment analysis

**Supplemental Table 13**: A comparison of nonsynonymous SNPs (benign + damaging) in top 1% ape diversity genes between Neandertal and modern human populations

